# Isolation of nucleic acids from low biomass samples: detection and removal of sRNA contaminants

**DOI:** 10.1101/232975

**Authors:** Anna Heintz-Buschart, Dilmurat Yusuf, Anne Kaysen, Alton Etheridge, Joëlle V. Fritz, Patrick May, Carine de Beaufort, Bimal B. Upadhyaya, Anubrata Ghosal, David J. Galas, Paul Wilmes

## Abstract

**Background:** Sequencing-based analyses of low-biomass samples are known to be prone to misinterpretation due to the potential presence of contaminating molecules derived from laboratory reagents and environments. Due to its inherent instability, contamination with RNA is usually considered to be unlikely.

**Results:** Here we report the presence of small RNA (sRNA) contaminants in widely used microRNA extraction kits and means for their depletion. Sequencing of sRNAs extracted from human plasma samples was performed and significant levels of non-human (exogenous) sequences were detected. The source of the most abundant of these sequences could be traced to the microRNA extraction columns by qPCR-based analysis of laboratory reagents. The presence of artefactual sequences originating from the confirmed contaminants were furthermore replicated in a range of published datasets. To avoid artefacts in future experiments, several protocols for the removal of the contaminants were elaborated, minimal amounts of starting material for artefact-free analyses were defined, and the reduction of contaminant levels for identification of *bona fide* sequences using ‘ultraclean’ extraction kits was confirmed.

**Conclusion:** This is the first report of the presence of RNA molecules as contaminants in laboratory reagents. The described protocols should be applied in the future to avoid confounding sRNA studies.

## BACKGROUND

The characterization of different classes of small RNAs (sRNAs) in tissues and bodily fluids holds great promise in understanding human physiology as well as in health-related applications. In blood plasma, microRNAs and other sRNAs are relatively stable, and microRNAs in particular are thought to reflect a system-wide state, making them potential biomarkers for a multitude of human diseases [1]. Different mechanisms of sRNA delivery as a means of long-distance intercellular communication have been recognized in several eukaryotes [2–7]. In addition, inter-individual, inter-species and even inter-kingdom communications via sRNAs have been proposed [8–12], and some cases of microRNA-based control by the host [13,14] or pathogens [15,16] have been demonstrated.

As exogenous RNAs have been detected in the blood plasma of humans and mice [17,18], the potential for exogenous RNA-based signalling in mammals is the subject of significant current debate [19,20]. Diet-derived exogenous microRNAs have been proposed to exert an influence on human physiology [21,22], as have bacterial RNAs, which can be secreted in the protective environment of outer membrane vesicles [23–25]. However, a heated discussion has at the same time been triggered around the genuineness of the observations of these exogenous sRNAs in human blood [26–28] and the possibility of dietary uptake of sRNAs [29–31]. This discussion happens at a time where DNA sequencing-based analyses of low-biomass samples have been recognized to be prone to confounding by contaminants [32]. From initial sample handling [33], to extraction kits [34], to sequencing reagents [35], multiple sources of DNA contamination and artefactual sequencing data have been described.

Here, we report the contamination of widely used silica-based columns for the isolation of micro- and other small RNAs with RNA, which was apparent from sRNA sequencing data and was subsequently validated by qPCR. These artefactual sRNA sequences were also apparent in numerous published datasets. Furthermore, approaches for the depletion of the contaminants from the columns as well as an evaluation of a newer ultra-clean kit are presented, along with the determination of a minimum safe input volume to suppress the signal of the contaminant sequences in RNA sequencing data of human blood plasma samples. The potential presence of *bona fide* exogenous sRNA species in human plasma is examined. Finally, recommendations for the control and interpretation of sRNA sequencing data from low-biomass samples are provided.

## RESULTS

### Initial detection of exogenous sRNAs in human blood plasma

sRNA was extracted from 100 μl blood plasma samples of ten healthy individuals and sequenced using regular RNeasy columns (workflow in **Figure 1**). The read profiles were mined for putative exogenous (non-human) sequences (Material and Methods). Among the potential exogenous sequences were 19 sequences that occurred with more than 1,000 counts per million (cpm) in all samples. To rule out sequencing errors or contamination during sequencing library preparation, a qPCR approach was developed to assess the presence of non-human sequences in the sRNA preparations from plasma. Six of the 19 highly abundant sRNA sequences from plasma that could not be mapped to the human genome were chosen for validation by qPCR (**Table 1**).

**Figure 1.**
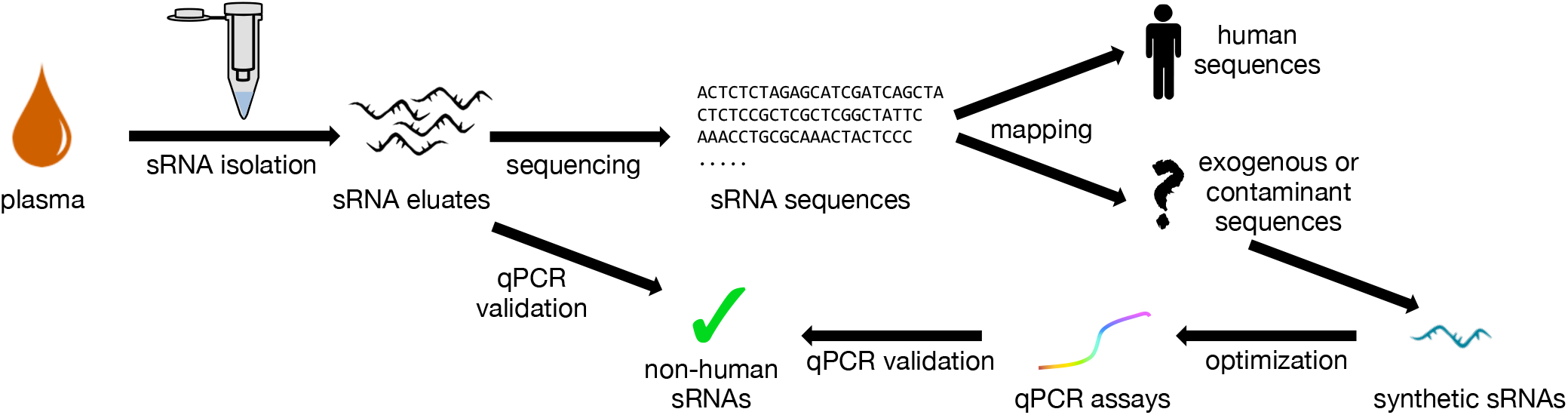
Workflow. Workflow of the initial screen for and validation of exogenous sRNA sequences in human plasma samples.

**Table 1.**
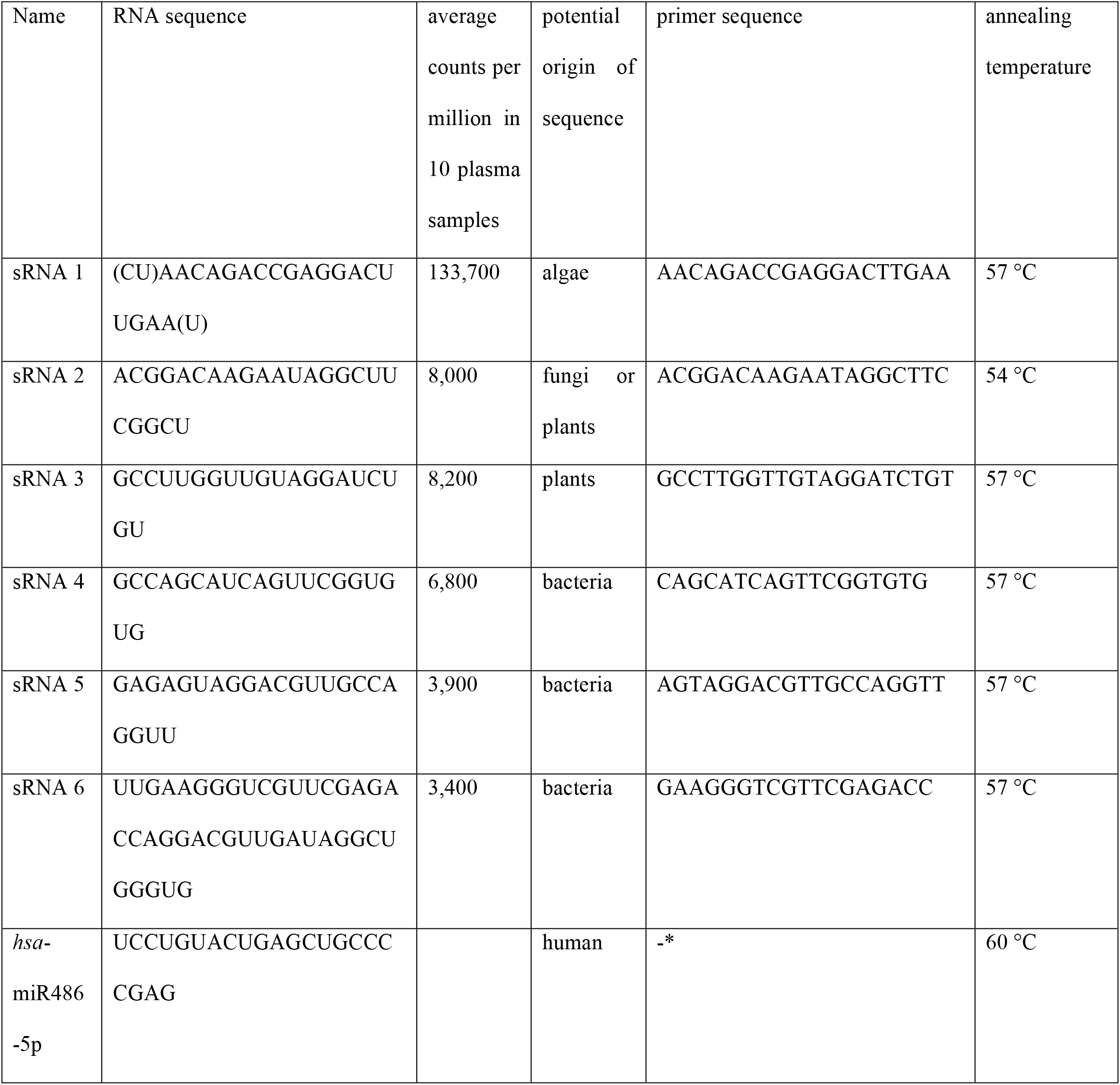
Sequences of non-human sRNAs found in plasma preparations, synthetic sRNA templates, primers and annealing temperatures.

### qPCR assays for putative exogenous sRNAs in human blood plasma

Synthetic sRNAs with the putative exogenous sequences found in plasma were poly-adenylated and reverse transcribed to yield cDNA, used for optimisation of PCR primers and conditions (**Table 1**). All primer sets yielded amplicons with single peaks in melting temperature analysis and efficiency values above 80 %. The optimised qPCR assays were then employed to test for the presence of the highly abundant sRNAs potentially representing exogenous sequences (workflow in **Figure 1**) in the human plasma samples used for the initial sequencing experiment. The qPCR assays confirmed the presence of these sRNAs in the sRNA preparations used for sequencing (**Figure 2A**), yielding amplicons with melting temperatures expected from the synthetic sRNAs. To rule out contamination of the water used in the sRNA preparations, a water control was also examined. No amplification was observed in all but one assay, where amplification of a product with a different melting temperature occurred (**Figure 2A**). Thus, for the assays, contamination of the water could be ruled out.

**Figure 2.**
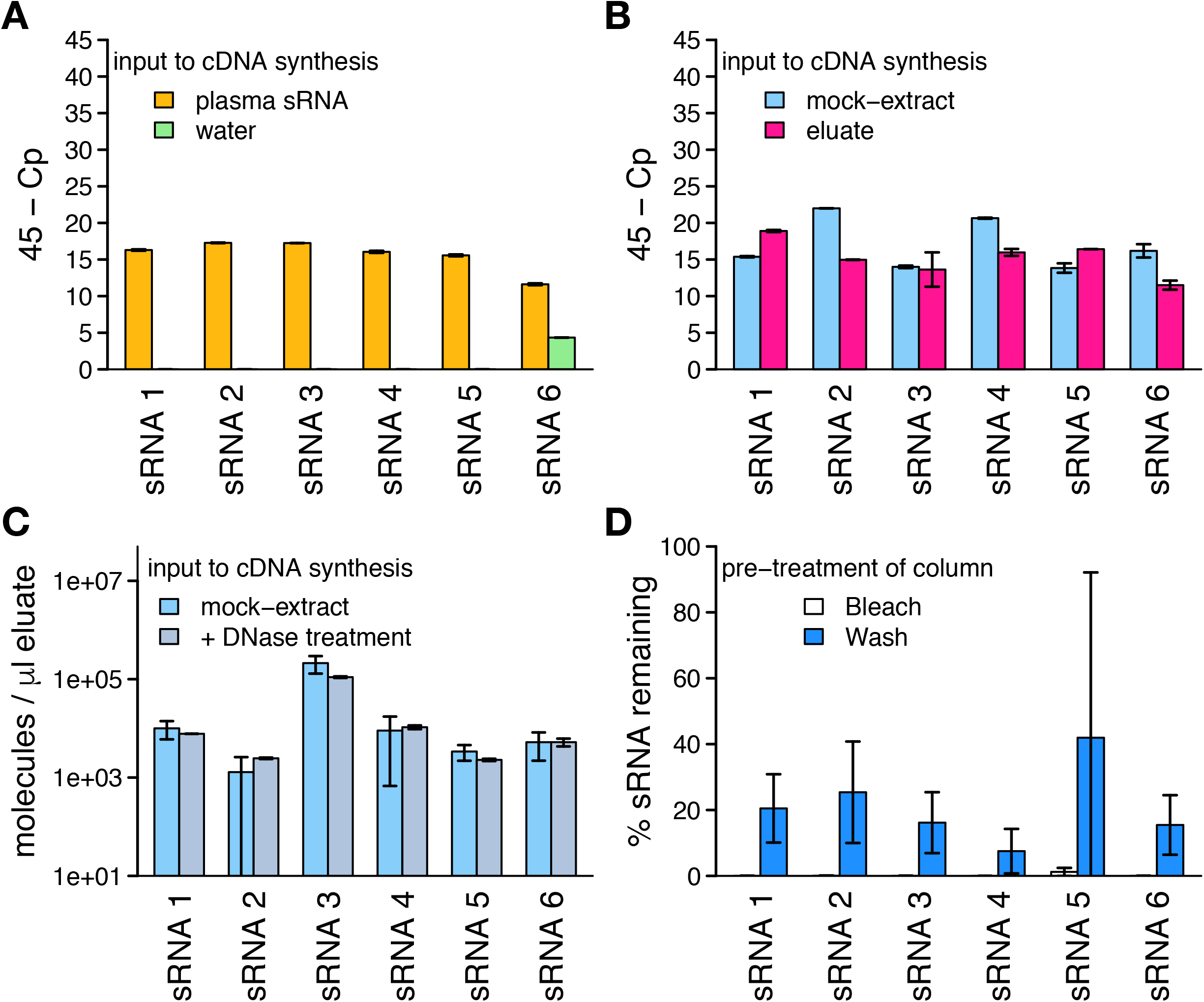
Detection of non-human sRNA species in column eluates and their removal from columns. **A**) qPCR amplification of six non-human sRNA species in extracts from human plasma and qPCR control (water). **B**) Detection of the same sRNA species in mock-extracts without input to extract columns and water passed through extraction columns (“eluate”). **C**) Levels of the same sRNA species in mock-extracts without and with DNase treatment during the extraction. **D**) Relative levels of sRNA remaining after pre-treatment of extraction columns with bleach or washing ten times with water, detected after eluting columns with water. **All**: mean results of three experiments, measured in reaction duplicates; error bars represent one standard deviation. Experiments displayed in panels **B** and **D** were performed on the same batch of columns, **A** and **C** on independent batches.

### Non-human sequences derived from column contaminants

To analyse whether the validated non-human sequences occurring in the sRNA extracts of plasma were present in any lab wear, a series of control experiments were carried out (**Additional Figure 1**). When nucleic acid- and RNase-free water (QIAGEN) was used as input to the miRNeasy Serum/Plasma kit (QIAGEN) instead of plasma (“mock-extraction”), all tested non-human sequences could be amplified from the mock-extract (**Figure 2B**). This indicates that one of the components of the extraction kit or lab-ware was contaminated with the non-human sequences. To locate the source of contamination, mock-extractions were performed by omitting single steps of the RNA-isolation protocol except for the elution step. Amplification from the resulting mock-extracts was tested for the most abundant non-human sequence (sRNA 1). In all cases, the sRNA 1 could be amplified (data not shown). We therefore carried out a simple experiment, in which nucleic acid- and RNase-free water was passed through an otherwise untreated spin column. From this column eluate, all target sequences could be amplified, in contrast to the nucleic acid- and RNase-free water (**Figure 2B**). The most abundant non-human sequences in the plasma sequencing experiments were therefore most likely contaminants originating from the untreated RNeasy columns.

### Detection of contaminant sequences in public datasets

To assess whether our observation of contaminant sRNAs was also pertinent in other sequencing datasets of low-input samples, the levels of confirmed contaminant sRNA sequences in published datasets [17,18,29,36–53] were assessed. Irrespective of the RNA isolation procedure applied, nontarget sequences were detected (making up between 5 and over 99 % of the sequencing libraries for the human samples; **Additional Table 1**). As shown in **Figure 3**, the six contaminant sequences which had been confirmed by qPCR were found in all analysed samples of low biomass samples which were extracted with regular miRNeasy kits, but the sequences were found at lower levels in studies with more biomass input [29,37,39] and hardly ever [40] in studies where samples were extracted using other methods (**Additional Table 1**). Within each study where the confirmed contaminant sequences were detected, the relative levels of the contaminant sequences were remarkably stable (**Additional Figure 2**).

**Figure 3.**
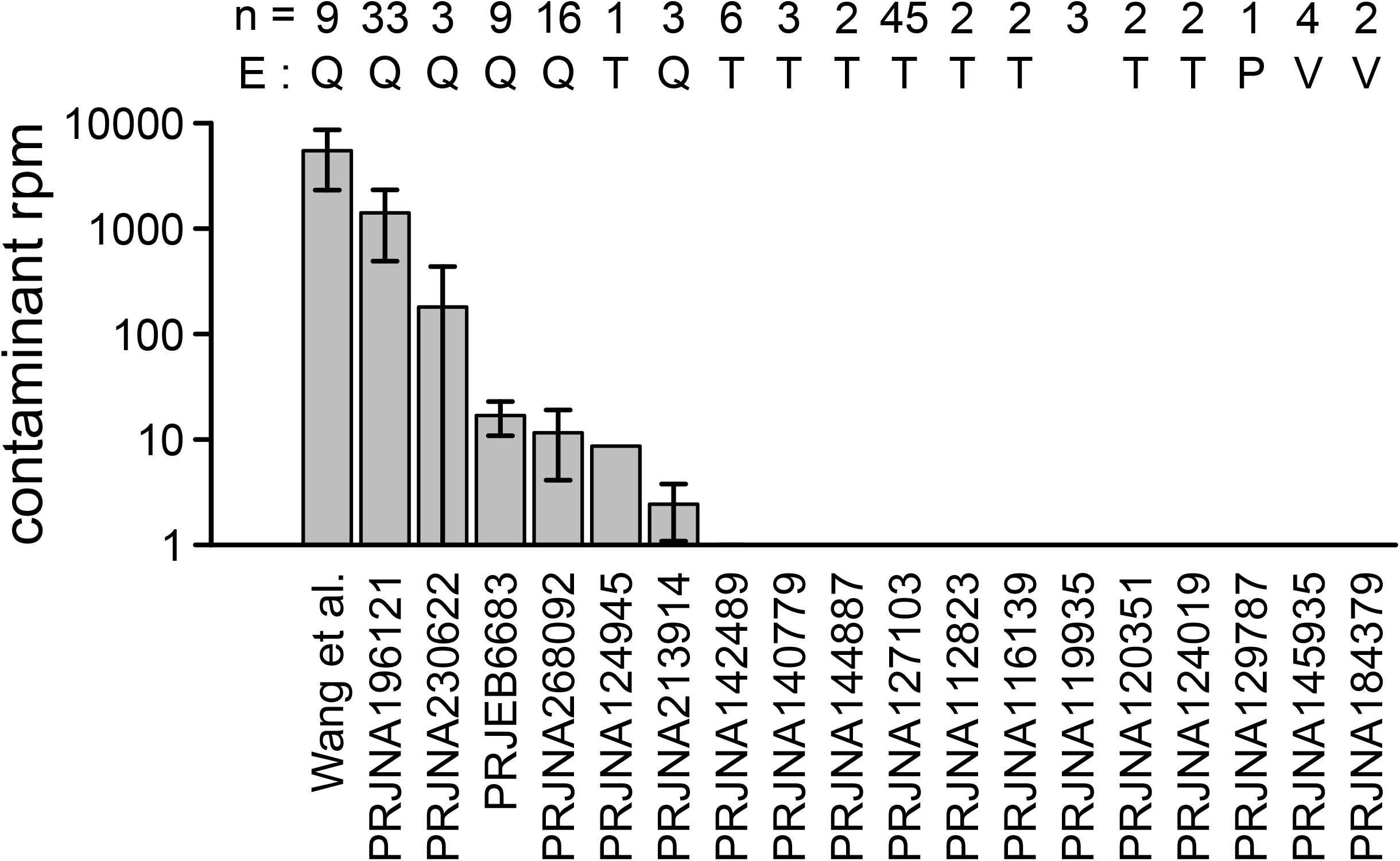
Detection of contaminant sequences in published sRNA sequencing datasets of low biomass samples. Datasets are referenced by NCBI bioproject accession or first author of the published manuscript. n: number of samples in the dataset. E: extraction kit used (if this information is available) – Q: regular miRNeasy (QIAGEN), T: TRIzol (Thermo Fisher), P: mirVana PARIS RNA extraction kit (Thermo Fisher), V: mirVana RNA extraction kit with phenol. rpm: reads per million. Error bars indicate one standard deviation.

### Depletion of contaminants from isolation columns

In order to eliminate contamination from the columns to allow their use in studies of environmental samples or potential exogenous sRNAs from human samples, we were interested in the nature of these contaminants. The fact that they can be poly-adenylated by RNA-poly-A-polymerase points to them being RNA. Treatment of the eluate with RNase prior to cDNA preparation also abolished amplification (data not shown), but on-column DNase digest did not reduce their levels (**Figure 2C**). These findings suggest that the contaminants were RNAs.

Contaminating sequences could potentially be removed from the RNeasy columns using RNase, but as RNases are notoriously difficult to inactivate and RNases remaining on the column would be detrimental to sRNA recovery, an alternative means of removing RNA was deemed desirable. Loading and incubation of RNeasy columns with the oxidant sodium hypochlorite and subsequent washing with RNase-free water to remove traces of the oxidant reduced amplifyability of unwanted sRNA by at least 100 times (**Figure 2D**), while retaining the columns’ efficiency to isolate sRNAs from samples applied afterwards. Elimination of contaminant sRNAs from the RNeasy columns by washing with RNase-free water (**Figure 2D**; average +/− standard deviation of the contaminant reduction by 80 +/− 10 %) or treatment with sodium hydroxide (average +/− standard deviation of the contaminant reduction by 70 +/− 15 %) was not sufficient to remove the contaminants completely.

### Ultra-clean extraction kits

Recently, RNeasy columns from an ultra-clean production have become available from QIAGEN within the miRNeasy Serum/Plasma Advanced Kit. We compared the levels of the previously analysed contaminant sequences in the flow-through of mock-extractions using 4 batches of ultraclean RNeasy columns to 2 batches of the regular columns by qPCR. In all cases, marked reductions in the contaminant levels were observed in the clean columns (**Figure 4A**; 4 to 4,000 fold; median 60). To obtain an overview over potential other contaminants, sRNA sequencing of the mock-extracts from these six batches of spin columns was performed. With regards to the six previously analysed contaminant sequences, the results were similar to those of the qPCR assays (**Additional Figure 3**). Additionally, for the ultra-clean RNeasy columns, a smaller spectrum of other potential contaminant sequences was observed (**Figure 4B&C**) and those sequences made up a smaller proportion of the eluate sequences (**Figure 4D**).

**Figure 4.**
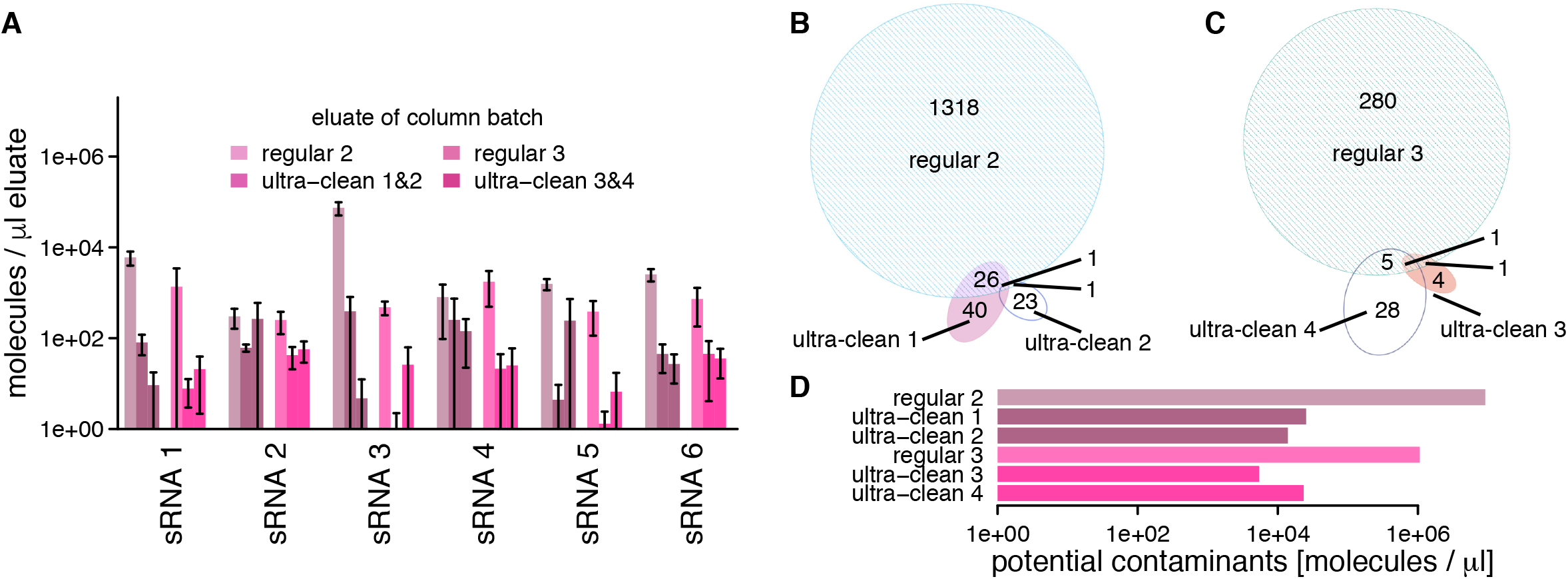
Confirmed and potential contaminant sequences in eluates of regular and ultra-clean RNeasy spin columns. **A**) Levels of contaminant sequences in eluates of two batches of regular and four batches of ultra-clean spin columns, based on qPCR; ultra-clean batches 1 and 2 are cleaned-up versions of regular batch 2 and ultra-clean batches 3 and 4 are cleaned-up versions of regular batch 3; error bars indicate one standard deviation. **B&C**) Numbers of different further potential contaminant sequences on the regular and ultra-clean spin columns from two different batches. **D**) Total levels of further potential contaminant sequences, based on sRNA sequencing data normalized to spike-in levels. cpm: counts per million.

As our initial analyses of plasma samples extracted using regular RNeasy spin columns had revealed contaminant levels of up to 7000 cpm, we were interested to define a safe input amount for human plasma for both column types that would be sufficient to suppress the contaminant signals to below 100 cpm. For this, we performed a titration experiment (**Additional Figure 3B**), isolating sRNA from a series of different input volumes of the same human plasma sample on four batches of RNeasy columns (2 batches of regular columns, 2 batches of ultra-clean columns) with subsequent sequencing. As expected from reagent contaminants, the observed levels of the contaminant sequences were generally inversely dependent on the plasma input volume (**Figure 5A**). In addition and in accordance with the earlier mock-extraction results, the levels of contaminant sequences were lower or they were completely absent in the ultra-clean columns (see levels for 100 μl input in **Figure 5B**). An input volume of 100 μl plasma was sufficient to reduce all contaminant sequences to below 100 cpm when using the ultra-clean spin columns.

**Figure 5.**
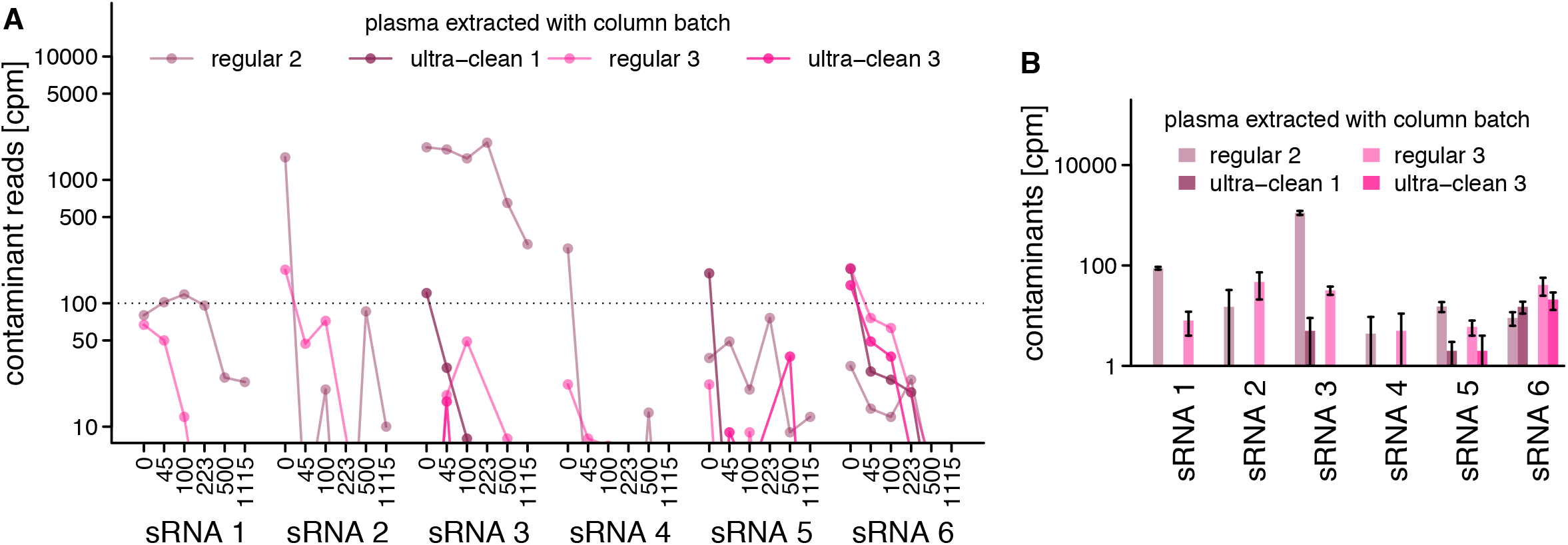
Titration experiment. Detection of contaminants in sRNA preparations of human plasma using different input volumes and extraction columns. **A**) Detected levels of the six contaminant sRNA sequences in sRNA sequencing data of preparations using 0 to 1115 μl human plasma and regular or ultra-clean RNeasy spin columns. **B**) Detailed view of the data displayed in A for 100 μl human plasma as input to regular and ultra-clean RNeasy spin columns. cpm: counts per million; error bars indicate one standard deviation.

### Potential plasma-derived exogenous RNAs

Finally, to detect potential exogenous sRNAs, we mined the plasma datasets used in the well-controlled titration experiment for sequences that do not originate from the human genome and were not detected in any of the mock-extracts. On average, 5 % of the sequencing reads of sRNA isolated from plasma did not map to the human genome. 127 sequences which did not map to the human genome assembly hg38 were detected in the majority of the plasma samples and were not represented in the control samples (empty libraries, column eluates or water). Out of these, 3 sequences had low complexity and 81 could be matched to sequences in the NCBI-nr that are not part of the current version of the human genome assembly (hg38) but annotated as human sequences or to sequences from other vertebrates. Of the 43 remaining sequences which matched to bacterial, fungal or plant sequences, 22 matched best to genera which have previously been identified as a source of contaminations of sequencing kits [35]. The remaining 21 sequences displayed very low (up to 47 cpm), yet consistent relative abundances in the 28 replicates of a plasma sample from the one healthy individual. Their potential origins were heterogeneous, including fungi and bacteria, with a notable enrichment in *Lactobacillus* sequences (**Additional Table 2**).

## DISCUSSION

Several instances of contamination of laboratory reagents with DNA, which can confound the analysis of sequencing data, have been reported in recent years [32,35,54,55]. In contrast, the contamination of reagents with RNA has not yet been reported. Contamination with RNA is usually considered very unlikely, due to the ubiquitous presence of RNases in the environment and RNA’s lower chemical stability due to being prone to hydrolysis, especially at higher pH. However, our results suggest that the detected contaminants were not DNA, but RNA, because treatment with RNase and not DNase could decrease the contaminant load. In addition, the contaminating molecules could not be amplified without poly-adenylation and reverse-transcription. The stability of the contaminants is likely due to the extraction columns being RNase-free and their silica protecting loaded sRNAs from degradation. While the results presented here focused on one manufacturer’s spin column-based extraction kit, for which contaminants were validated, other RNA-stabilizing or extraction reagents may carry RNA contaminations. This is suggested by previously observed significant batch effects of sequencing data derived from samples extracted with a number of different extraction kits [27]. Based on the analysis of the published data sets, where significant numbers of sequences that did not map to the source organism’s genome were found independent of the RNA extraction kit used, the potential contaminants in other extraction kit would have different sequences than the ones confirmed by qPCR here.

The results presented here should help to assess the question whether exogenous sRNA species derived from oral intake [18] or the human microbiome [17,38,56] really occur frequently in human plasma or are merely artefacts [26]. While the limited data from this study (one healthy person) points to very low levels and a small spectrum of potential foreign sRNAs, properly controlled studies using laboratory materials without contaminants on individuals or animals with conditions that limit gastrointestinal barrier function will shed more light on this important research question in the future.

## CONCLUSIONS

The reported contaminant sequences can confound studies of organisms whose transcriptomes contain sequences similar to the contaminants. They can also give rise to misinterpretation in studies without *a priori* knowledge of the present organisms as well as lead to the overestimation of miRNA yields in low-biomass samples. Therefore, based on the present study, care has to be taken when analysing low-input samples, in particular for surveys of environmental or otherwise undefined sources of RNAs. A number of recommendations can be conceived based on the presented data (**Figure 6**): Extraction columns should be obtained as clean as possible. Simple clean-up procedures can also reduce contaminants. The input mass of sRNA should be as high as possible, e.g. for human plasma volumes above 100 μl are preferable. Extraction controls should always be sequenced with the study samples. To facilitate library preparation for the extraction controls, spike-in RNAs with defined sequences can be used. They should be applied at concentrations similar to the levels of RNA found in the study samples. As the spike-in signal can drown out the contaminants, it is necessary to avoid too high concentrations for the spike-ins. Sequences found in the extraction controls should be treated as artefacts and removed from the sequencing data. Independent techniques that are more robust to low input material, such as qPCR or ddPCR, should be applied to both study samples and controls in case of doubt.

**Figure 6.**
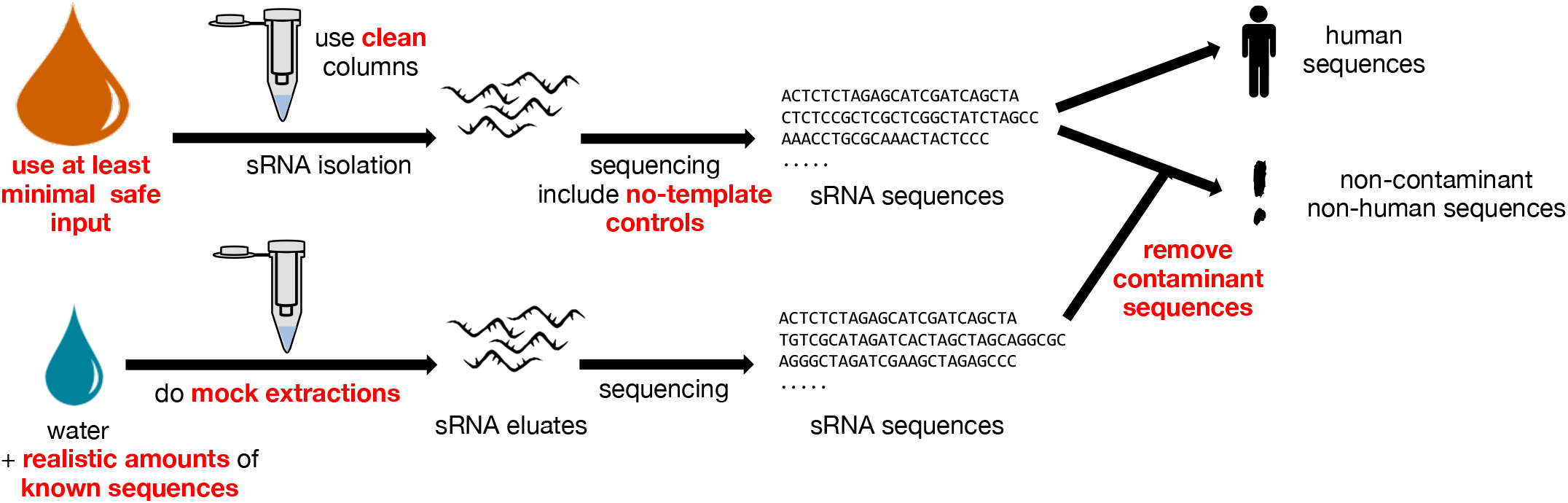
Summary. Recommendations for artefact-free analysis of sRNA by sequencing.

## METHODS

### Blood plasma sampling

Written informed consent was obtained from all blood donors. The sample collection and analysis was approved by the Comité d’Ethique de Recherche (CNER; Reference: 201110/05) and the National Commission for Data Protection in Luxembourg. Blood was collected by venepuncture into EDTA-treated tubes. Plasma was prepared immediately after blood collection by centrifugation (10 min at 1,000 × *g*) and platelets were depleted by a second centrifugation step (5 min at 10,000 × *g*). The blood plasma was flash-frozen in liquid nitrogen and stored at −80 °C until extraction.

### Use of sRNA isolation columns

Unless stated otherwise, 100 μl blood plasma was lysed using the QIAzol (QIAGEN) lysis reagent prior to binding to the column, as recommended by the manufacturer. RNeasy MinElute spin columns from the miRNeasy Serum/Plasma Kit (QIAGEN) were then loaded, washed and dried, and RNA was eluted as recommended by the manufacturer’s manual. We further tested four batches of ultra-clean RNeasy MinElute columns, which underwent an ultra-clean production process (UCP) to remove potential nucleic acid contaminations, including environmental sRNAs. These columns were treated as recommended in the manual of the miRNeasy Serum/Plasma Advanced Kit (QIAGEN). All eluates were stored at −80 °C until analysis.

For the mock-extractions, ultra-clean or regular RNeasy columns were loaded with the aqueous phase from a QIAzol extraction of nucleic acid- and RNase-free water (QIAGEN) instead of plasma. For mock-extractions with a defined spike-in, the aqueous phase was spiked with synthetic *hsa*-miR-486-3p RNA (Eurogentec) to yield 40,000 copies per μl eluate. To obtain column eluates, spin columns were not loaded, washed or dried. Instead, 14 μl of RNase-free water (QIAGEN) was applied directly to a new column and centrifuged for 1 min.

To eliminate environmental sRNAs from the regular RNeasy columns, the columns were incubated with 500 μl of a sodium hypochlorite solution (Sigma; diluted in nuclease free water (Invitrogen) to approx. 0.5 %) for 10 min at room temperature. Columns were subsequently washed 10 times with 500 μl nuclease free water (Invitrogen), before use. Similarly, in the attempt to remove sRNAs by application of sodium hydroxide, 500 μl 50 mM NaOH were incubated on the spin columns for 5 min, followed by incubation with 50 mM HCl for 5 min, prior to washing the columns 10 times with 500 μl nuclease-free water (Invitrogen) before use.

### Real-time PCR

5 μl of eluted RNA was polyadenylated and reverse-transcribed to cDNA using the qScript microRNA cDNA Synthesis Kit (Quanta BIOSCIENCES). 1 μl of cDNA (except for the initial plasma experiment, where 0.2 μl cDNA were used) was amplified by use of sequence-specific forward primers (see **Table 1**, obtained from Eurogentec) or the miR486-5p specific assay from Quanta BIOSCIENCES, PerfeCTa Universal PCR Primer and PerfeCTa SYBR Green SuperMix (Quanta BIOSCIENCES) in a total reaction volume of 10 μl. Primers were added at a final concentration of 0.2 μM. Primer design and amplification settings were optimised with respect to reaction efficiency and specificity. Efficiency was calculated using a dilution series covering seven orders of magnitude of template cDNA reverse transcribed from synthetic sRNA. Real-time PCR was performed on a LightCycler^®^ 480 Real-Time PCR System (Roche) including denaturation at 95 °C for 2 min and 40 cycles of 95 °C for 5 sec, 54–60 °C for 15 sec (for annealing temperatures see **Table 1**), and 72 °C for 15 sec. All reactions were carried out in duplicates. No-template-controls were performed analogously with water as input. Cp values were obtained using the second derivative procedure provided by the LightCycler^®^ 480 Software, Version 1.5. Cp data were analysed using the comparative C_T_ method (ΔΔCT).

### sRNA seq: library preparation and sequencing

sRNA libraries were made using the TruSeq small RNA library preparation kit (Illumina) according to the manufacturer’s instructions, except that the 3’ and 5’ adapters were diluted 1:3 before use. PCR-amplified libraries were size selected using a PippinHT instrument (Sage Science), collecting the range of 121–163 bp. Completed, size-selected libraries were run on a High Sensitivity DNA chip on a 2100 Bioanalyzer (Agilent) to assess library quality. Concentration was determined by qPCR using the NEBNext Library Quant kit (NEB). Libraries were pooled, diluted and sequenced with 75 cycle single-end reads on a NextSeq 500 (Illumina) according the manufacturer’s instructions. The sequencing reads can be accessed at NCBI’s short read archive via PRJNA419919 (for sample identifiers and accessions see **Additional Table 1**).

### Initial analysis: plasma-derived sRNA sequencing data

For the initial analysis of plasma-derived sRNA sequencing data, FastQC (http://www.bioinformatics.babraham.ac.uk/projects/fastqc) was used to determine over-represented primer and adapter sequences, which were subsequently removed using cutadapt (http://dx.doi.org/10.14806/ej.17.1.200). This step was repeated recursively until no over-represented primer or adapter sequences were detected. 5’-Ns were removed using fastx_clipper of the FASTX-toolkit. Trimmed reads were quality-filtered using fastq_quality_filter of the FASTX-toolkit (with −q 30 −p 90; http://hannonlab.cshl.edu/fastx_toolkit). Finally, identical reads were collapsed, retaining the read abundance information using fastx_collapser of the FASTX-toolkit. The collapsed reads were mapped against the human genome (GRCh37), including RefSeq exon junction sequences, as well as prokaryotic, viral, fungal, plant and animal genomes from Genbank [57] and the Human Microbiome Project [58] using Novoalign V2.08.02 (http://www.novocraft.com; **Additional Table 3**). These organisms were selected based on the presence in the human microbiome, human nutrition and the public availability of the genomes. As reads were commonly mapping to genomic sequences of multiple organisms, and random alignment can easily occur between short sequences and reference genomes, the following approach was taken to refine their taxonomic classification: First, reads were attributed to the human genome if they mapped to it. Secondly, reads mapping to each reference genome was compared to mapping of a shuffled decoy read set. Based on this, the list of reference genomes was limited to the genomes recruiting at least one read with a minimum length of 25 nt. Loci on non-human genomes were established by the position of the mapping reads. The number of mapping reads per locus was adjusted using a previously established cross-mapping correction [59]. Finally, the sequences of the loci, the number of mapping reads and their potential taxonomy were extracted.

### sRNA sequence analysis of controls

For the subsequent analysis of the mock-extractions, column eluates and nucleic acid- and RNase-free water, and no-template controls as well as human plasma samples, extracted using either regular or ultra-clean RNeasy columns, the trimming and quality check of the reads was done analogously to the description above. Collapsed reads were mapped against the most recent version of the human genome (hg38) either to remove operator-derived sequences or to distinguish the reads mapping to the human genome in the different datasets. Sequencing was performed in two batches, with one batch filling an entire flow cell, and one mixed with other samples. The latter batch of samples was sequenced on the same flow cell as sRNAs extracted from *Salmonella typhimurium* LT2. To avoid misinterpretations due to multiplexing errors, reads mapping to *Salmonella typhimurium* LT2 [60] (Genbank accession AE006468) were additionally removed in this batch. To limit the analysis to only frequently occurring sequences and therefore avoid over-interpretation of erroneous sequences, only read sequences that were found at least 30 times in all analysed samples together were retained for further analysis. Public sRNA datasets of low-input samples (see **Additional Table 1**) were analysed in a fashion analogous to the study’s control and plasma samples. As the published studies consisted of different numbers of samples, no overall threshold was imposed, but to limit the analysis to frequently occurring sequences, singleton reads were removed.

To compare the sequencing results to the qPCR-based results and to detect the same sequences in public datasets, reads matching the sequences assayed by qPCR were determined by clustering the trimmed, filtered and collapsed sRNA reads with 100 % sequence identity and 14 nt alignment length with the primer sequences, while allowing the sRNA reads to be longer than the primer sequences, using CD-HIT-EST-2D (parameters -c 1 -n 8 -G 0 -A 14 -S2 40 -g 1 -r 0) [61].

To compare the diversity and levels of putative contaminant sequences in the different samples, identical reads derived from all study samples (that did not map to the human genome) were clustered using CD-HIT-EST [61], and a table with the number of reads sequenced for each sample per sequence was created using R v.3.0.2. This table was also used to extract candidate sequences from the study plasma samples that are likely exogenous plasma sRNAs, based on the following criteria: for a sequence to be considered a potential exogenous plasma sRNA, it had to be non-identical to any of the sequences assigned to the confirmed contaminant sequences (**Table 1**), and it had to be absent from at least 90 % of the controls (no-library controls, water and spike-in controls, eluates and mock-extracts) and never detected in any of these controls with at least 10 copy numbers, and it had to be detected by more than 3 reads in more than 7 of the 28 libraries generated from the plasma titration experiment. These thresholds were chosen in order to make the analysis robust against multiplexing errors (e.g. which would result in false-negative identifications if a sequence that is very dominant in a plasma sample is falsely assigned to the control-samples), while at the same time making it sensitive to low-abundant sequences (which would not be detected in every library). To confirm the non-human origin and find potential microbial taxa of origin for these sequences, they were subsequently searched within the NCBI nr database using megablast and blastn web tools, with parameters auto-set for short inputs [62–64]. All sequences with best hits to human sequences or other vertebrates were removed, because they were potentially human. The remaining sequences were matched against a set of genera previously reported [35] to be common sequencing kit contaminants. Sequences with better hits to non-contaminant taxa than contaminant taxa were kept as potential exogenous sequences.

### Additional Files

The following Additional Files are available online: **Additional Figures 1–3; Additional Table 1**: list of the generated datasets and analysed published datasets; **Additional Table 2**: potential exogenous sRNA sequences detected in human plasma after removal of contaminants; **Additional Table 3**: list of the species whose reference genomes and cDNA collections were used in the initial analysis.

#### LIST OF ABBREVIATIONS

qPRC: : real-time quantitative polymerase chain reaction
sRNA: : small RNA

## DECLARATIONS

### Ethics approval and consent to participate

Written informed consent was obtained from all blood donors. The sample collection and analysis was approved by the Comité d’Ethique de Recherche (CNER; Reference: 201110/05) and the National Commission for Data Protection in Luxembourg.

### Consent for publication

Written consent for analysis of genetic material and publication was obtained from all blood donors.

### Availability of data and materials

The datasets generated and analysed during the current study are available in the NCBI short read archive under BioProject PRJNA419919. Human reads from some datasets generated and analysed during the current study are not publicly available due to privacy concerns, but are available from the corresponding authors on reasonable request. Accessions of publically available data analysed during the current study are listed in **Additional Table 1**. Scripts for the analysis of the data from sRNA sequencing of column eluates and the plasma titration experiment is available at https://git.ufz.de/metaOmics/contaminomics.

### Authors ‘ contributions

AH-B designed the experiments, performed experiments and sequencing data analyses, coordinated the study and wrote the manuscript. DY designed and performed the initial sequencing data analyses. AK, JVF and AG performed experiments. AE performed the sRNA sequencing. PM and BBU performed additional computational analyses. CdB obtained donor consents, performed the blood sampling and contributed to the initiation of the study. DJG and PW initiated and supervised the study. DY, AK, AE, JVF, PM and PW contributed to the writing of the manuscript. All authors contributed to the interpretation of the data and read and approved the final manuscript.

## Acknowledgements

*In silico* analyses presented in this paper were carried out using the HPC facilities of the University of Luxembourg [65] whose administrators are acknowledged for excellent support.

## REFERENCES

1. Mitchell PS, Parkin RK, Kroh EM, Fritz BR, Wyman SK, Pogosova-Agadjanyan EL, et al. Circulating microRNAs as stable blood-based markers for cancer detection. Proc. Natl. Acad. Sci. U.S.A. 2008;105:10513–8.

2. Valadi H, Ekström K, Bossios A, Sjöstrand M, Lee JJ, Lötvall JO. Exosome-mediated transfer of mRNAs and microRNAs is a novel mechanism of genetic exchange between cells. Nat. Cell Biol. 2007;9:654–9.

3. Zernecke A, Bidzhekov K, Noels H, Shagdarsuren E, Gan L, Denecke B, et al. Delivery of microRNA-126 by apoptotic bodies induces CXCL12-dependent vascular protection. Sci Signal. 2009;2:ra81.

4. Pegtel DM, Cosmopoulos K, Thorley-Lawson DA, van Eijndhoven MAJ, Hopmans ES, Lindenberg JL, et al. Functional delivery of viral miRNAs via exosomes. Proc. Natl. Acad. Sci. U.S.A. 2010;107:6328–33.

5. Molnar A, Melnyk CW, Bassett A, Hardcastle TJ, Dunn R, Baulcombe DC. Small silencing RNAs in plants are mobile and direct epigenetic modification in recipient cells. Science. 2010;328:872–5.

6. Vickers KC, Palmisano BT, Shoucri BM, Shamburek RD, Remaley AT. MicroRNAs are transported in plasma and delivered to recipient cells by high-density lipoproteins. Nat. Cell Biol. Nature Publishing Group; 2011;13:423–33.

7. Arroyo JD, Chevillet JR, Kroh EM, Ruf IK, Pritchard CC, Gibson DF, et al. Argonaute2 complexes carry a population of circulating microRNAs independent of vesicles in human plasma. Proc. Natl. Acad. Sci. U.S.A. 2011;108:5003–8.

8. Tomilov AA, Tomilova NB, Wroblewski T, Michelmore R, Yoder JI. Trans-specific gene silencing between host and parasitic plants. Plant J. 2008;56:389–97.

9. Kosaka N, Izumi H, Sekine K, Ochiya T. microRNA as a new immune-regulatory agent in breast milk. Silence. 2010;1:7.

10. Knip M, Constantin ME, Thordal-Christensen H. Trans-kingdom cross-talk: small RNAs on the move. PLoS Genet. 2014;10:e1004602.

11. Fritz JV, Heintz-Buschart A, Ghosal A, Wampach L, Etheridge A, Galas D, et al. Sources and Functions of Extracellular Small RNAs in Human Circulation. Annu. Rev. Nutr. 2016;36:301–36.

12. Koeppen K, Hampton TH, Jarek M, Scharfe M, Gerber SA, Mielcarz DW, et al. A Novel Mechanism of Host-Pathogen Interaction through sRNA in Bacterial Outer Membrane Vesicles. PLoS Pathog. 2016;12:e1005672.

13. LaMonte G, Philip N, Reardon J, Lacsina JR, Majoros W, Chapman L, et al. Translocation of sickle cell erythrocyte microRNAs into Plasmodium falciparum inhibits parasite translation and contributes to malaria resistance. Cell Host Microbe. 2012;12:187–99.

14. Liu S, da Cunha AP, Rezende RM, Cialic R, Wei Z, Bry L, et al. The Host Shapes the Gut Microbiota via Fecal MicroRNA. Cell Host Microbe. 2016;19:32–43.

15. Weiberg A, Wang M, Lin F-M, Zhao H, Zhang Z, Kaloshian I, et al. Fungal small RNAs suppress plant immunity by hijacking host RNA interference pathways. Science. 2013;342:118–23.

16. Buck AH, Coakley G, Simbari F, McSorley HJ, Quintana JF, Le Bihan T, et al. Exosomes secreted by nematode parasites transfer small RNAs to mammalian cells and modulate innate immunity. Nature Communications. 2014;5:5488.

17. Wang K, Li H, Yuan Y, Etheridge A, Zhou Y, Huang D, et al. The complex exogenous RNA spectra in human plasma: an interface with human gut biota? Wang K, Li H, Yuan Y, Etheridge A, Zhou Y, Huang D, et al., editors. PLoS ONE. 2012;7:e51009.

18. Zhang Y, Wiggins BE, Lawrence C, Petrick J, Ivashuta S, Heck G. Analysis of plant-derived miRNAs in animal small RNA datasets. BMC Genomics. 2012;13:381.

19. Zhang L, Hou D, Chen X, Li D, Zhu L, Zhang Y, et al. Exogenous plant MIR168a specifically targets mammalian LDLRAP1: evidence of cross-kingdom regulation by microRNA. Nature Publishing Group. 2011;22:107–26.

20. Zhou Z, Li X, Liu J, Dong L, Chen Q, Liu J, et al. Honeysuckle-encoded atypical microRNA2911 directly targets influenza A viruses. Cell Research. 2015;25:39–49.

21. Liang G, Zhu Y, Sun B, Shao Y, Jing A, Wang J, et al. Assessing the survival of exogenous plant microRNA in mice. Food Sci Nutr. 2014;2:380–8.

22. Baier SR, Nguyen C, Xie F, Wood JR, Zempleni J. MicroRNAs Are Absorbed in Biologically Meaningful Amounts from Nutritionally Relevant Doses of Cow Milk and Affect Gene Expression in Peripheral Blood Mononuclear Cells, HEK-293 Kidney Cell Cultures, and Mouse Livers. Journal of Nutrition. 2014;144:1495–500.

23. Ghosal A, Upadhyaya BB, Fritz JV, Heintz-Buschart A, Desai MS, Yusuf D, et al. The extracellular RNA complement of Escherichia coli. MicrobiologyOpen. 2015;4:252–66.

24. Celluzzi A, Masotti A. How Our Other Genome Controls Our Epi-Genome. Trends Microbiol. 2016;24:777–87.

25. Blenkiron C, Simonov D, Muthukaruppan A, Tsai P, Dauros P, Green S, et al. Uropathogenic Escherichia coli Releases Extracellular Vesicles That Are Associated with RNA. Cascales E, editor. PLoS ONE. 2016;11:e0160440–16.

26. Witwer KW. Contamination or artifacts may explain reports of plant miRNAs in humans. The Journal of Nutritional Biochemistry. 2015;26:1685.

27. Kang W, Bang-Berthelsen CH, Holm A, Houben AJS, Müller AH, Thymann T, et al. Survey of 800+ data sets from human tissue and body fluid reveals xenomiRs are likely artifacts. RNA. Cold Spring Harbor Lab; 2017;23:433–45.

28. Witwer KW, Zhang C-Y. Diet-derived microRNAs: unicorn or silver bullet? Genes & Nutrition. BioMed Central; 2017;12:15.

29. Dickinson B, Zhang Y, Petrick JS, Heck G, Ivashuta S, Marshall WS. Lack of detectable oral bioavailability of plant microRNAs after feeding in mice. Nat. Biotechnol. 2013;31:965–7.

30. Witwer KW, Hirschi KD. Transfer and functional consequences of dietary microRNAs in vertebrates: Concepts in search of corroboration. Bioessays. 2014;36:394–406.

31. Title AC, Denzler R, Stoffel M. Uptake and Function Studies of Maternal Milk-derived MicroRNAs. Journal of Biological Chemistry. American Society for Biochemistry and Molecular Biology; 2015;290:23680–91.

32. Lusk RW. Diverse and widespread contamination evident in the unmapped depths of high throughput sequencing data. PLoS ONE. 2014;9:e110808.

33. Salzberg SL, Breitwieser FP, Kumar A, Hao H, Burger P, Rodriguez FJ, et al. Next-generation sequencing in neuropathologic diagnosis of infections of the nervous system. Neurol Neuroimmunol Neuroinflamm. 2016;3:e251.

34. Naccache SN, Greninger AL, Lee D, Coffey LL, Phan T, Rein-Weston A, et al. The Perils of Pathogen Discovery: Origin of a Novel Parvovirus-Like Hybrid Genome Traced to Nucleic Acid Extraction Spin Columns. J. Virol. 2013;87:11966–77.

35. Salter SJ, Cox MJ, Turek EM, Calus ST, Cookson WO, Moffatt MF, et al. Reagent and laboratory contamination can critically impact sequence-based microbiome analyses. BMC Biol. BioMed Central; 2014;12:87.

36. Huang X, Yuan T, Tschannen M, Sun Z, Jacob H, Du M, et al. Characterization of human plasma-derived exosomal RNAs by deep sequencing. BMC Genomics. 2013;14:319.

37. Spornraft M, Kirchner B, Haase B, Benes V, Pfaffl MW, Riedmaier I. Optimization of Extraction of Circulating RNAs from Plasma – Enabling Small RNA Sequencing. Antoniewski C, editor. PLoS ONE. 2014;9:e107259.

38. Beatty M, Guduric-Fuchs J, Brown E, Bridgett S, Chakravarthy U, Hogg RE, et al. Small RNAs from plants, bacteria and fungi within the order Hypocreales are ubiquitous in human plasma. BMC Genomics. BioMed Central; 2014;15:933.

39. Santa-Maria I, Alaniz ME, Renwick N, Cela C, Fulga TA, Van Vactor D, et al. Dysregulation of microRNA-219 promotes neurodegeneration through post-transcriptional regulation of tau. J. Clin. Invest. 2015;125:681–6.

40. Taft RJ, Simons C, Nahkuri S, Oey H, Korbie DJ, Mercer TR, et al. Nuclear-localized tiny RNAs are associated with transcription initiation and splice sites in metazoans. Nat Struct Mol Biol. 2010;17:1030–4.

41. Chen C, Ai H, Ren J, Li W, Li P, Qiao R, et al. A global view of porcine transcriptome in three tissues from a full-sib pair with extreme phenotypes in growth and fat deposition by paired-end RNA sequencing. BMC Genomics. BioMed Central; 2011;12:448.

42. Liu J-L, Liang X-H, Su R-W, Lei W, Jia B, Feng X-H, et al. Combined Analysis of MicroRNome and 3′-UTRome Reveals a Species-specific Regulation of Progesterone Receptor Expression in the Endometrium of Rhesus Monkey. J. Biol. Chem. 2012;287:13899–910.

43. Lebedeva S, Jens M, Theil K, Schwanhäusser B, Selbach M, Landthaler M, et al. Transcriptome-wide Analysis of Regulatory Interactions of the RNA-Binding Protein HuR. Mol. Cell. Elsevier Inc; 2011;43:340–52.

44. Kuchen S, Resch W, Yamane A, Kuo N, Li Z, Chakraborty T, et al. Regulation of MicroRNA Expression and Abundance during Lymphopoiesis. Immunity. Elsevier Ltd; 2010;32:828–39.

45. Wei Y, Chen S, Yang P, Ma Z, Kang L. Characterization and comparative profiling of the small RNA transcriptomes in two phases of locust. Genome Biology. 2009;10:R6.

46. Mayr C, Bartel DP. Widespread Shortening of 3′ UTRs by Alternative Cleavage and Polyadenylation Activates Oncogenes in Cancer Cells. Cell. Elsevier Ltd; 2009;138:673–84.

47. Su R-W, Lei W, Liu J-L, Zhang Z-R, Jia B, Feng X-H, et al. The Integrative Analysis of microRNA and mRNA Expression in Mouse Uterus under Delayed Implantation and Activation. Wang H, editor. PLoS ONE. 2010;5:e15513–8.

48. Chen X, Yu X, Cai Y, Zheng H, Yu D, Liu G, et al. Next-generation small RNA sequencing for microRNAs profiling in the honey bee Apis mellifera. Insect Mol Biol. 2010;19:799–805.

49. Legeai F, Rizk G, Walsh T, Edwards O, Gordon K, Lavenier D, et al. Bioinformatic prediction, deep sequencing of microRNAs and expression analysis during phenotypic plasticity in the pea aphid, Acyrthosiphon pisum. BMC Genomics. BioMed Central; 2010;11:281.

50. Vaz C, Ahmad HM, Sharma P, Gupta R, Kumar L, Kulshreshtha R, et al. Analysis of microRNA transcriptome by deep sequencing of small RNA libraries of peripheral blood. BMC Genomics. BioMed Central; 2010;11:288.

51. Liu S, Li D, Li Q, Zhao P, Xiang Z, Xia Q. MicroRNAs of Bombyx mori identified by Solexa sequencing. BMC Genomics. BioMed Central; 2010;11:148.

52. Lian L, Qu L, Chen Y, Lamont SJ, Yang N. A Systematic Analysis of miRNA Transcriptome in Marek’s Disease Virus-Induced Lymphoma Reveals Novel and Differentially Expressed miRNAs. Watson M, editor. PLoS ONE. 2012;7:e51003–13.

53. Nolte-‘t Hoen ENM, Buermans HPJ, Waasdorp M, Stoorvogel W, Wauben MHM, ‘t Hoen PAC. Deep sequencing of RNA from immune cell-derived vesicles uncovers the selective incorporation of small non-coding RNA biotypes with potential regulatory functions. Nucleic Acids Research. 2012;40:9272–85.

54. Lauder AP, Roche AM, Sherrill-Mix S, Bailey A, Laughlin AL, Bittinger K, et al. Comparison of placenta samples with contamination controls does not provide evidence for a distinct placenta microbiota. Microbiome. BioMed Central; 2016;4:29.

55. Glassing A, Dowd SE, Galandiuk S, Davis B, Chiodini RJ. Inherent bacterial DNA contamination of extraction and sequencing reagents may affect interpretation of microbiota in low bacterial biomass samples. Gut Pathog. BioMed Central; 2016;8:24.

56. Yeri A, Courtright A, Reiman R, Carlson E, Beecroft T, Janss A, et al. Total Extracellular Small RNA Profiles from Plasma, Saliva, and Urine of Healthy Subjects. Sci Rep. 2017;7:44061.

57. Benson DA, Cavanaugh M, Clark K, Karsch-Mizrachi I, Lipman DJ, Ostell J, et al. GenBank. Nucleic Acids Res. 2012;41:D36–D42.

58. The NIH HMP Working Group, Peterson J, Garges S, Giovanni M, McInnes P, Wang L, et al. The NIH Human Microbiome Project. Genome Research. 2009;19:2317–23.

59. de Hoon MJL, Taft RJ, Hashimoto T, Kanamori-Katayama M, Kawaji H, Kawano M, et al. Cross-mapping and the identification of editing sites in mature microRNAs in high-throughput sequencing libraries. Genome Research. 2010;20:257–64.

60. McClelland M, Sanderson KE, Spieth J, Clifton SW, Latreille P, Courtney L, et al. Complete genome sequence of Salmonella enterica serovar Typhimurium LT2. Nature. 2001;413:852–6.

61. Fu L, Niu B, Zhu Z, Wu S, Li W. CD-HIT: accelerated for clustering the next-generation sequencing data. Bioinformatics. 2012;28:3150–2.

62. Altschul SF, Gish W, Miller W, Myers EW, Lipman DJ. Basic local alignment search tool. J. Mol. Biol. 1990;215:403–10.

63. Morgulis A, Coulouris G, Raytselis Y, Madden TL, Agarwala R, Schäffer AA. Database indexing for production MegaBLAST searches. Bioinformatics. 2008;24:1757–64.

64. Database resources of the National Center for Biotechnology Information. Nucleic Acids Res. 2016;44:D7–D19.

65. Varrette S, Bouvry P, Cartiaux H, Georgatos F. Management of an academic HPC cluster: The UL experience. IEEE; 2014;:959–67.

